# Molecular and clinical analysis of 27 German patients with Leber congenital amaurosis

**DOI:** 10.1101/428177

**Authors:** Nicole Weisschuh, Britta Feldhaus, Muhammad Imran Khan, Frans P. M. Cremers, Susanne Kohl, Bernd Wissinger, Ditta Zobor

## Abstract

Leber congenital amaurosis (LCA) is the earliest and most severe form of all inherited retinal dystrophies (IRD) and the most frequent cause of inherited blindness in children. The phenotypic overlap with other early-onset and severe IRDs as well as difficulties associated with the ophthalmic examination of infants can complicate the clinical diagnosis. To date, 25 genes have been implicated in the pathogenesis of LCA. The disorder is usually inherited in an autosomal recessive fashion, although rare dominant cases have been reported. We report the mutation spectra and frequency of genes in 27 German index patients initially diagnosed with LCA. A total of 108 LCA- and other genes implicated in IRD were analysed using a cost-effective targeted next-generation sequencing procedure based on molecular inversion probes (MIPs). Sequencing and variant filtering led to the identification of putative pathogenic variants in 25 cases, thereby leading to a detection rate of 93%. The mutation spectrum comprises 34 different alleles, 17 of which are novel. In line with previous studies, the genetic results led to a revision of the initial clinical diagnosis in a substantial proportion of cases, demonstrating the importance of genetic testing in IRD. In addition, our detection rate of 93% shows that MIPs are a cost-efficient and sensitive tool for targeted next-generation sequencing in IRD.

## Introduction

Leber congenital amaurosis (LCA, MIM #204000) was first described by Theodor Leber in 1869 and refers to a heterogeneous group of severe, mostly recessively inherited, early infantile-onset retinal dystrophies with typically extinguished electroretinograms (ERGs). Later, a separate group of milder disease phenotypes, with some preservation of the ERG responses, the so-called “early-onset severe retinal dystrophy” (EOSRD) or “severe early childhood onset retinal dystrophy” has been described. LCA and EOSRD together are the most severe and earliest forms of all inherited retinal diseases (IRDs). They affect 20% of blind children and account for 5% of all IRDs [1]. In Germany, the estimated number of cases is 2000 (source: Pro Retina Deutschland e. V.). To date, mutations in 25 genes have been associated with LCA (https://sph.uth.edu/retnet/). A substantial proportion of cases (10-20%) remain unsolved despite extensive molecular testing [2-4]. This is due to technical limitations as copy number variations often remain undetected in datasets derived from capture panels or whole exome sequencing, but also because of the focus on coding regions in most diagnostic settings which will not detect deep intronic variants acting on splicing or variants in regulatory sequences.

There is a considerable clinical and genetic overlap between LCA, EOSRD and other types of IRD, therefore, an accurate clinical diagnosis cannot always be made at the first visit of the young patients. Furthermore, the clinical examination of infants is challenging or limited. Hence, the initial clinical diagnosis sometimes has to be revised once genetic results are available.

For a long time the genetic heterogeneity of LCA (and IRD in general) hampered DNA-based (molecular) diagnoses, since parallel screening of all associated genes requires next generation sequencing approaches, for which reimbursement to the patient is often not guaranteed. We sought for a cost-effective and sensitive approach to obtain a molecular diagnosis for 27 patients that had been diagnosed with LCA at the University Eye Hospital Tuebingen. The present study focuses on these genetically unsolved cases, which were screened for sequence variants in 108 genes associated with non-syndromic IRD by a cost-effective targeted panel-based next-generation sequencing approach.

## Materials and Methods

### Subjects and clinical assessment

In this study we included 27 unrelated patients of German origin with a clinical diagnosis of LCA who were not genetically pre-investigated. Their clinical diagnosis was established by standard clinical ophthalmologic examinations including patient history, psychophysical and electrophysiological examinations. Genomic DNA of patients was extracted from peripheral blood using standard protocols. Samples from all patients and family members were recruited in accordance with the principles of the Declaration of Helsinki and were obtained with written informed consent accompanying the patients’ samples. The study was approved by the institutional review board of the Ethics Committee of the University Hospital of Tuebingen.

### Sequencing analysis

Molecular testing was performed by targeted next-generation sequencing at a core facility (Department of Human Genetics, Radboud University Nijmegen Medical Centre). We used molecular inversion probes (MIPs) with 5-bp molecular tags to conduct targeted next generation sequencing of 108 genes associated with IRD (see S1 Table). The 1,524 coding exons and the 10 bp flanking each exon were targeted with 6,129 probes for an overall target size of 647,574 bp. On average, 4-6 MIPs cover one exon. The panel also includes the frequent LCA-associated pathogenic intronic variant c.2992+1655A>G in *CEP290* [5]. Pooled and phosphorylated probes were added to the capture reactions with 100 ng of genomic DNA from each individual to produce a library for each individual. The libraries were amplified with 21 cycles of PCR, during which an 8-bp sample barcode was introduced. The barcoded libraries were then pooled and purified with AMPureXP beads (Beckman-Coulter). Sequencing was performed on an Illumina NextSeq 500 system. Demultiplexed BAM files were aligned to a human reference sequence (UCSC Genome Browser hg19) via the Burrows-Wheeler Aligner (BWA) v.0.6.2 [6]. In-house automated data analysis pipeline and variant interpretation tools were used for variant calling. Rare and potentially disease-causing variants were confirmed by Sanger sequencing using standard protocols. Sanger sequencing was also used to screen for the recurrent c.2843G>A/p.C948Y variant in the *CRB1* gene.

### Variant filtering and classification

Only non-synonymous single nucleotide variants (nsSNVs), nonsense variants, putative splice site (±10 bps) variants, insertions, duplications and deletions represented by more than 20 sequence reads were considered for further analysis. In addition, variants with a minor allele frequency (MAF) >0.5% in the Genome Aggregation Database (gnomAD) Version r2.0.2 [7] were excluded from further investigation. For variant classification we applied the terminology proposed by the American College of Medical Genetics and Genomics and the Association for Molecular Pathology [8].

### *In silico* predictions

The potential pathogenicity of the missense changes identified in this study was assessed using four online prediction software tools, namely SIFT (http://sift.bii.a-star.edu.sg/) [9], PolyPhen-2 (http://genetics.bwh.harvard.edu/pph2/) [10], Mutation Taster (www.mutationtaster.org/) [11], and Provean (http://provean.jcvi.org/) [12].

## Results

Utilizing our capture panel technology, we were able to obtain an average of 1.2 million reads on target per sample, with an average coverage of 213 reads per probe. Moreover, an average of 88% of targeted regions had 10x coverage or more, which was sufficient for accurate variant calling. The pipeline initially called an average of 532 single nucleotide variants and 64 insertions/deletions for each sample. Putative pathogenic variants were identified in 25/27 index cases (Table 1), thereby achieving a detection rate of 93%. All putative disease-associated variants were validated by conventional Sanger sequencing. Homozygosity was observed for eight patients (26%): variants were seen in true homozygous state in four patients and in apparent homozygous state in four patients, respectively. Two patients were hemizygous, and compound heterozygosity was observed for four patients based on the analysis of paternal alleles. *Trans* configuration of variants could not be demonstrated for 11 patients because DNA of family members was not available and the respective variants were located too far apart for allelic cloning. In patient 26, a single heterozygous variant in *IMPG2* was observed. In patient 27, no putative disease-causing variants were identified. The mutation spectrum comprises 34 different alleles, 17 of which are novel. All novel variants were deposited to the ClinVar database (https://www.ncbi.nlm.nih.gov/clinvar/) [13] with accession codes provided in Table 1.

**Table 1.** Putative pathogenic variants in 25 unrelated German patients initially diagnosed with LCA

| Patient Nr. | Final diagnosis | Gene | Allele 1 | Reference | ClinVar accession no. | ACMG category | Allele 2 | Reference | ClinVar accession no. | ACMG category | Segregation performed |
| --- | --- | --- | --- | --- | --- | --- | --- | --- | --- | --- | --- |
| **Solved by putative pathogenic mutations in known LCA genes** |  |  |  |  |  |  |  |  |  |  |  |
| 1 | LCA | *AIPL1* | c.857A>T/p.D286V | this study | pending | VUS | c.857A>T/p.D286V | this study | pending | VUS | yes |
| 2 | LCA | *AIPL1* | c.834G>A/p.W278* | PMID: 10615133 | SCV000086966.1 | LP | c.277+6T>C/p.? | this study | pending | VUS | no |
| 3 | LCA | *CEP290* | c.2991+1655A>G/p.[C998*, =] | PMID: 16909394<br>PMID: 27151457 | SCV000021550.2 | LP | c.2991+1655A>G/p.[C998*, =] | PMID: 16909394<br>PMID: 27151457 | SCV000021550.2 | LP | yes |
| 4 | EOSRD | *CRB1* | c.2798G>A/p.C933Y | this study | pending | VUS | c.2843G>A/p.C948Y | PMID: 10508521 | SCV000056582.2 | VUS | yes |
| 5 | LCA | *CRB1* | c.4039del/p.T1347Lfs*5 | this study | pending | LP | c.2843G>A/p.C948Y | PMID: 10508521 | SCV000056582.2 | VUS | no |
| 6 | LCA | *CRB1* | c.410del/p.P137Lfs*11 | this study | pending | LP | c.2843G>A/p.C948Y | PMID: 10508521 | SCV000056582.2 | VUS | no |
| 7 | LCA | *CRB1* | c.70+1G>A/p.? | this study | pending | LP | c.2042G>A/p.C681Y | PMID: 11231775 | SCV000118458.1 | VUS | yes |
| 8 | EOSRD | *CRB1* | c.2308G>A/p.G770S | PMID: 27113771 | SCV000282584.1 | VUS | c.2843G>A/p.C948Y | PMID: 10508521 | SCV000056582.2 | VUS | no |
| 9 | LCA | *CRB1* | c.2072G>A/p.W691* | this study | pending | LP | c.2843G>A/p.C948Y | PMID: 10508521 | SCV000056582.2 | VUS | no |
| 10 | LCA | *NMNAT1* | c.12dup/p.E5Rfs*4 | PMID: 24940029 | n.a. | LP | c.769G>A/p.E257K | PMID: 22842231 | SCV000053426.1 | VUS | no |
| 11 | EOSRD | *RD3* | c.180C>A/p.Y60* | PMID: 22531706 | SCV000222653.1 | LP | c.180C>A/p.Y60* | PMID: 22531706 | SCV000222653.1 | LP | no |
| 12 | LCA | *RPE65* | c.110G>C/p.W37S | this study | pending | VUS | c.722A>G/p.H241R | this study | pending | VUS | no |
| 13 | LCA | *RPE65* | c.203A>C/p.H68P | this study | pending | VUS | c.825C>G/p.Y275* | this study | pending | LP | no |
| 14 | LCA | *RPGRIP1* | c.2440C>T/p.R814* | this study | pending | LP | c.2440C>T/p.R814* | this study | pending | LP | no |
| 15 | LCA | *RPGRIP1* | c.1303A>T/p.K435* | PMID: 27208204 | SCV000282616.1 | LP | c.801-25_c.843del | this study | pending | LP | yes |
| 16 | LCA | *RPGRIP1* | c.2941C>T/p.R981* | PMID: 28041643 | SCV000599101.1 | LP | c.2941C>T/p.R981* | PMID: 28041643 | SCV000599101.1 | LP | no |
| 17 | LCA | *RPGRIP1* | c.800+1G>A/p.? | PMID: 16123401 | n.a. | LP | c.2718dup/p.N907* | PMID: 28714225 | n.a. | LP | yes |
| **Solved by putative pathogenic mutations in IRD genes not typically associated with LCA** |  |  |  |  |  |  |  |  |  |  |  |
| 18 | CRD | *ABCA4* | c.1765del/p.W589Gfs*60 | this study | pending | LP | c.1765del/p.W589Gfs*60 | this study | pending | LP | no |
| 19 | CRD | *ABCA4* | c.5461-10T>C/p.[T1821Vfs*13, T1821Dfs*6] | PMID: 15614537<br>PMID: 26976702 | SCV000028574.2 | LP | c.3377T>C/p.L1126P | PMID: 25066811 | SCV000281867.2 | VUS | no |
| 20 | CRD | *ABCA4* | c.3259G>A/E1087K | PMID: 9054934 | SCV000598963.1 | VUS | c.5917del/p.V1973* | PMID: 10958763 | SCV000599004.1 | LP | no |
| 21 | CRD | *ABCA4* | c.5917del/p.V1973* | PMID: 10958763 | SCV000599004.1 | LP | c.5917del/p.V1973* | PMID: 10958763 | SCV000599004.1 | LP | yes |
| 22 | CSNB | *CACNA1F* | c.4504C>T/p.R1502* | this study | pending | LP | - |  |  |  | no |
| 23 | CRD | *CDHR1* | c.634G>A/p.A212T | PMID: 16288196 | n.a. | VUS | c.1132C>T/p.R378W | this study | pending | VUS | no |
| 24 | RP | *PROM1* | c.1209_1229/p.Q403_S410delinsH | PMID: 24265693 | SCV000575406.4 | VUS | c.1209_1229/p.Q403_S410delinsH | PMID: 24265693 | SCV000575406.4 | VUS | yes |
| 25 | XRP | *RP2* | c.314G>A/p.C105Y | this study | pending | VUS | - |  |  |  | no |

LCA, Leber congenital amaurosis; EOSRD, early-onset severe retinal dystrophy; CRD, cone-rod dystrophy; CSNB, congenital stationary nightblindness; RP, retinitis pigmentosa; XRP, X-linked RP; VUS, variant of uncertain significance; LP, likely pathogenic; n.a., not available.

The variants comprise 14 missense variants, eight nonsense variants, seven deletions or duplications leading to a frame-shift, three canonical splice site variants, two non-canonical splice site variants and one in-frame deletion. Pathogenicity was interpreted in accordance with the American College of Medical Genetics guidelines [8]. The respective categories are given in Table 1. Missense variants that have never been reported before were analysed using different *in silico* prediction algorithms. These scores, together with the MAFs sourced from the gnomAD browser are shown in Table 2.

**Table 2.** Assessment of pathogenicity of missense variants identified in this study

|  |  |  |  |  |  |  |  |  |
| --- | --- | --- | --- | --- | --- | --- | --- | --- |
| Gene | Variant | gnomAD MAF | Mutation Taster | Polyphen | SIFT | Provean | phyloP | Grantham Score |
| **ABCA4** | c.3377T>C/p.L1126P | 4.061e-6 | Disease causing (0.99) | Probably damaging (1.0) | Damaging (0.0) | Deleterious (-6.51) | 3.60 | 98 |
| **ABCA4** | c.3259G>A/E1087K | 1.624e-5 | Disease causing (0.99) | Probably damaging (1.0) | Damaging (0.0) | Deleterious (-3.84) | 6.22 | 56 |
| **AIPL1** | c.857A>T/p.D286V | none | Disease causing (0.99) | Probably damaging (1.0) | Damaging (0.0) | Deleterious (-8.31) | 4.09 | 152 |
| **CDHR1** | c.634G>A/p.A212T | 0.0001312 | Disease causing (0.99) | Probably damaging (0.99) | Damaging (0.0) | Deleterious (-3.22) | 4.93 | 58 |
| **CDHR1** | c.1132C>T/p.R378W | 7.584e-5 | Disease causing (0.99) | Probably damaging (1.0) | Damaging (0.02) | Deleterious (-3.84) | 1.25 | 101 |
| **CRB1** | c.2798G>A/p.C933Y | none | Disease causing (0.99) | Probably damaging (0.99) | Damaging (0.0) | Deleterious (-9.66) | 5.69 | 194 |
| **CRB1** | c.2308G>A/p.G770S | 2.036e-5 | Disease causing (0.99) | Probably damaging (1.0) | Tolerated (0.06) | Deleterious (-5.48) | 5.69 | 56 |
| **CRB1** | c.2843G>A/p.C948Y | 0.0002027 | Disease causing (0.99) | Probably damaging (0.99) | Damaging (0.0) | Deleterious (-9.66) | 5.31 | 194 |
| **CRB1** | c.2042G>A/p.C681Y | 4.067e-6 | Disease causing (0.99) | Probably damaging (1.0) | Damaging (0.0) | Deleterious (-10.74) | 5.74 | 194 |
| **NMNAT1** | c.769G>A/p.E257K | 0.0006968 | Disease causing (0.99) | Benign (0.09) | Tolerated (0.52) | Neutral (-2.31) | 3.87 | 56 |
| **RP2** | c.314G>A/p.C105Y | none | Disease causing (0.99) | Probably damaging (1.0) | Damaging (0.0) | Deleterious (-8.6) | 5.50 | 194 |
| **RPE65** | c.110G>C/p.W37S | none | Disease causing (0.99) | Probably damaging (1.0) | Damaging (0.02) | Deleterious (-12.62) | 5.78 | 177 |
| **RPE65** | c.722A>G/p.H241R | none | Disease causing (0.99) | Probably damaging (1.0) | Damaging (0.0) | Deleterious (-7.58) | 4.74 | 29 |
| **RPE65** | c.203A>C/p.H68P | none | Disease causing (0.99) | Probably damaging (1.0) | Tolerated (0.06) | Deleterious (-9.34) | 4.78 | 77 |

MAF, minor allele frequency.

### LCA / EOSRD patients

A summary of clinical findings is shown in Table 3 including all 27 index patients. In 19 of 27 patients, the initial diagnosis of LCA/EOSRD was confirmed by the molecular genetic analysis. In all of these cases, disease onset was typically at birth or within the first months of life. Nystagmus and strabismus were common features, indicating the lack of visual development. Visual acuity was severely reduced in all cases, ranging from 0.2 (decimal) to no light perception (NLP). Where visual field testing was possible, only small residual visual islands could be detected. Fullfield ERGs were extinguished in each case at time of recording. Morphological findings included typical salt & pepper pigmentary changes of the retina, pale optic disks and attenuated retinal vessels. Patients showed a progressive disease history with severe visual impairment from the beginning. In the following, the LCA-associated genes that were found to be mutated in these patients are listed in detail.

**Table 3.**
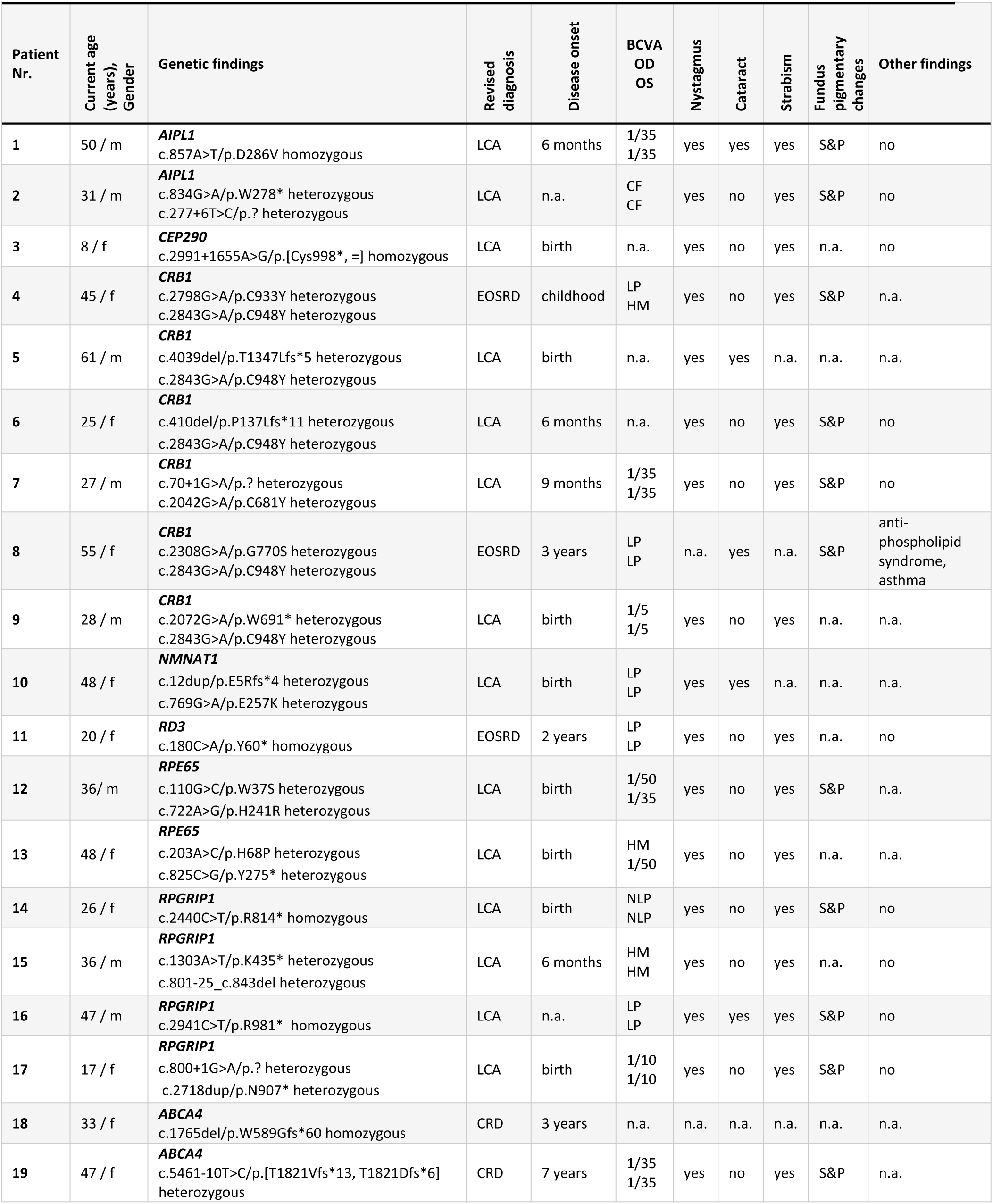
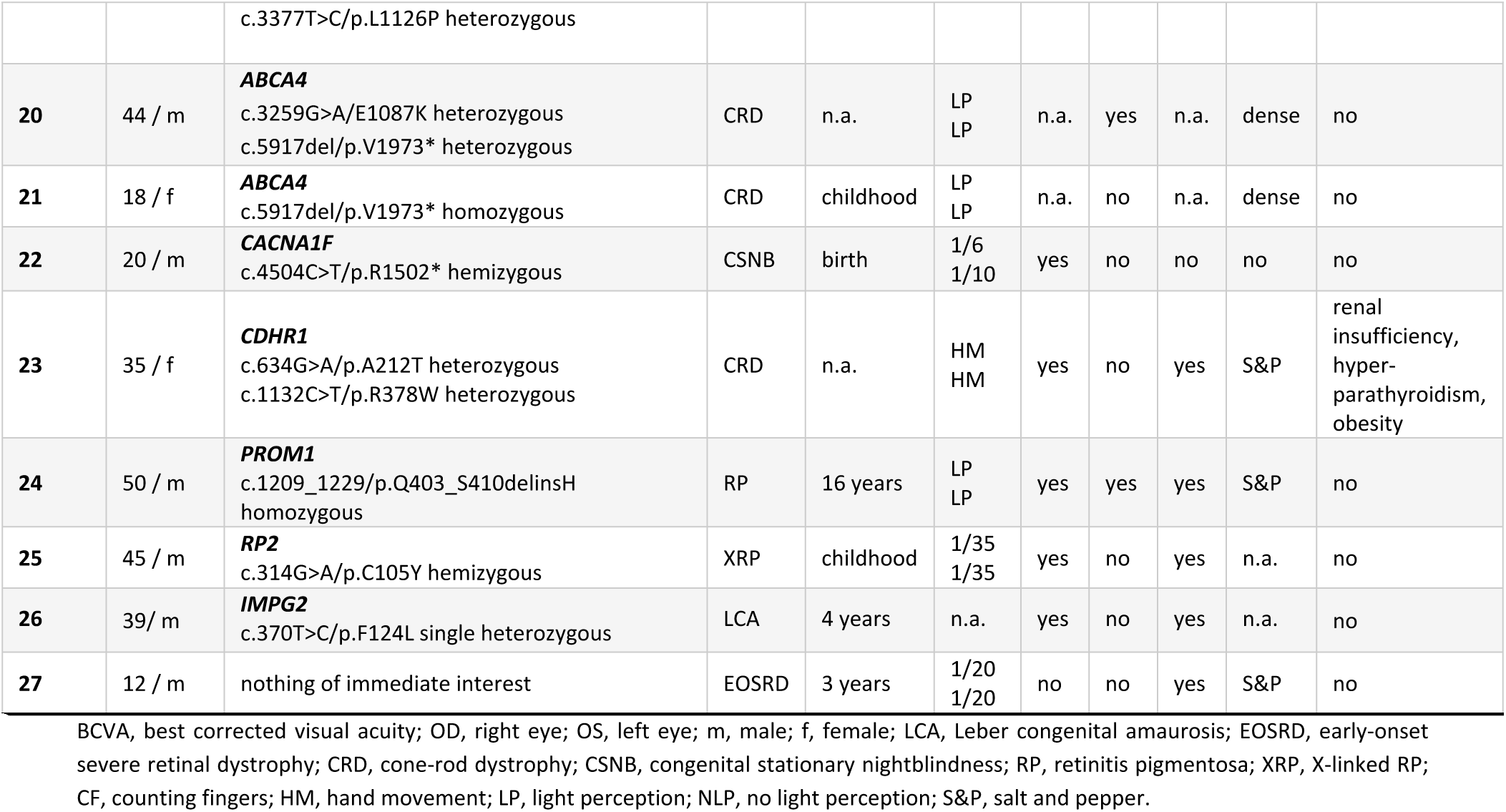
Summary of clinical findings

| Patient Nr. | Current age (years), Gender | Genetic findings | Revised diagnosis | Disease onset | BCVA OD OS | Nystagmus | Cataract | Strabism | Fundus pigmentary changes | Other findings |
| --- | --- | --- | --- | --- | --- | --- | --- | --- | --- | --- |
| 1 | 50 / m | **AIPL1**<br>c.857A>T/p.D286V homozygous | LCA | 6 months | 1/35<br>1/35 | yes | yes | yes | S&P | no |
| 2 | 31 / m | **AIPL1**<br>c.834G>A/p.W278* heterozygous<br>c.277+6T>C/p.? heterozygous | LCA | n.a. | CF<br>CF | yes | no | yes | S&P | no |
| 3 | 8 / f | **CEP290**<br>c.2991+1655A>G/p.[Cys998*, =] homozygous | LCA | birth | n.a. | yes | no | yes | n.a. | no |
| 4 | 45 / f | **CRB1**<br>c.2798G>A/p.C933Y heterozygous<br>c.2843G>A/p.C948Y heterozygous | EOSRD | childhood | LP<br>HM | yes | no | yes | S&P | n.a. |
| 5 | 61 / m | **CRB1**<br>c.4039del/p.T1347Lfs*5 heterozygous<br>c.2843G>A/p.C948Y heterozygous | LCA | birth | n.a. | yes | yes | n.a. | n.a. | n.a. |
| 6 | 25 / f | **CRB1**<br>c.410del/p.P137Lfs*11 heterozygous<br>c.2843G>A/p.C948Y heterozygous | LCA | 6 months | n.a. | yes | no | yes | S&P | no |
| 7 | 27 / m | **CRB1**<br>c.70+1G>A/p.? heterozygous<br>c.2042G>A/p.C681Y heterozygous | LCA | 9 months | 1/35<br>1/35 | yes | no | yes | S&P | no |
| 8 | 55 / f | **CRB1**<br>c.2308G>A/p.G770S heterozygous<br>c.2843G>A/p.C948Y heterozygous | EOSRD | 3 years | LP<br>LP | n.a. | yes | n.a. | S&P | anti-phospholipid syndrome, asthma |
| 9 | 28 / m | **CRB1**<br>c.2072G>A/p.W691* heterozygous<br>c.2843G>A/p.C948Y heterozygous | LCA | birth | 1/5<br>1/5 | yes | no | yes | n.a. | n.a. |
| 10 | 48 / f | **NMNAT1**<br>c.12dup/p.E5Rfs*4 heterozygous<br>c.769G>A/p.E257K heterozygous | LCA | birth | LP<br>LP | yes | yes | n.a. | n.a. | n.a. |
| 11 | 20 / f | **RD3**<br>c.180C>A/p.Y60* homozygous | EOSRD | 2 years | LP<br>LP | yes | no | yes | n.a. | no |
| 12 | 36 / m | **RPE65**<br>c.110G>C/p.W37S heterozygous<br>c.722A>G/p.H241R heterozygous | LCA | birth | 1/50<br>1/35 | yes | no | yes | S&P | n.a. |
| 13 | 48 / f | **RPE65**<br>c.203A>C/p.H68P heterozygous<br>c.825C>G/p.Y275* heterozygous | LCA | birth | HM<br>1/50 | yes | no | yes | n.a. | n.a. |
| 14 | 26 / f | **RPGRIP1**<br>c.2440C>T/p.R814* homozygous | LCA | birth | NLP<br>NLP | yes | no | yes | S&P | no |
| 15 | 36 / m | **RPGRIP1**<br>c.1303A>T/p.K435* heterozygous<br>c.801-25_c.843del heterozygous | LCA | 6 months | HM<br>HM | yes | no | yes | n.a. | no |
| 16 | 47 / m | **RPGRIP1**<br>c.2941C>T/p.R981* homozygous | LCA | n.a. | LP<br>LP | yes | yes | yes | S&P | no |
| 17 | 17 / f | **RPGRIP1**<br>c.800+1G>A/p.? heterozygous<br>c.2718dup/p.N907* heterozygous | LCA | birth | 1/10<br>1/10 | yes | no | yes | S&P | no |
| 18 | 33 / f | **ABCA4**<br>c.1765del/p.W589Gfs*60 homozygous | CRD | 3 years | n.a. | n.a. | n.a. | n.a. | n.a. | n.a. |
| 19 | 47 / f | **ABCA4**<br>c.5461-10T>C/p.[T1821Vfs*13, T1821Dfs*6] heterozygous c.3377T>C/p.L1126P heterozygous | CRD | 7 years | 1/35<br>1/35 | yes | no | yes | S&P | n.a. |
| 20 | 44 / m | **ABCA4**<br>c.3259G>A/E1087K heterozygous<br>c.5917del/p.V1973* heterozygous | CRD | n.a. | LP<br>LP | n.a. | yes | n.a. | dense | no |
| 21 | 18 / f | **ABCA4**<br>c.5917del/p.V1973* homozygous | CRD | childhood | LP<br>LP | n.a. | no | n.a. | dense | no |
| 22 | 20 / m | **CACNA1F**<br>c.4504C>T/p.R1502* hemizygous | CSNB | birth | 1/6<br>1/10 | yes | no | no | no | no |
| 23 | 35 / f | **CDHR1**<br>c.634G>A/p.A212T heterozygous<br>c.1132C>T/p.R378W heterozygous | CRD | n.a. | HM<br>HM | yes | no | yes | S&P | renal<br>insufficiency,<br>hyper-<br>parathyroidism,<br>obesity |
| 24 | 50 / m | **PROM1**<br>c.1209_1229/p.Q403_S410delinsH<br>homozygous | RP | 16 years | LP<br>LP | yes | yes | yes | S&P | no |
| 25 | 45 / m | **RP2**<br>c.314G>A/p.C105Y hemizygous | XRP | childhood | 1/35<br>1/35 | yes | no | yes | n.a. | no |
| 26 | 39 / m | **IMPG2**<br>c.370T>C/p.F124L single heterozygous | LCA | 4 years | n.a. | yes | no | yes | n.a. | no |
| 27 | 12 / m | nothing of immediate interest | EOSRD | 3 years | 1/20<br>1/20 | no | no | yes | S&P | no |

BCVA, best corrected visual acuity; OD, right eye; OS, left eye; m, male; f, female; LCA, Leber congenital amaurosis; EOSRD, early-onset severe retinal dystrophy; CRD, cone-rod dystrophy; CSNB, congenital stationary nightblindness; RP, retinitis pigmentosa; XRP, X-linked RP; CF, counting fingers; HM, hand movement; LP, light perception; NLP, no light perception; S&P, salt and pepper.

### CRB1

*CRB1* variants were detected in six patients (22.2%; 6/27). In total, eight variants were identified, including one novel nonsense, two novel frame-shifting deletions, one novel canonical splice site variant and one novel missense variant. Compound heterozygosity could only be demonstrated in two patients. Five patients were heterozygous for the recurrent c.2843G>A/p.C948Y variant, which has been reported to represent 23–31% of all *CRB1* disease-associated alleles [14-15]. Of note, this particular variant was not covered by the MIPs in our assay, but we screened all patients by conventional Sanger Sequencing for this variant, because of its known high frequency and relevance (MAF 0.0002027).

### RPGRIP1

Of the six potentially disease-causing variants in *RPGRIP1* detected in four patients (14.8%; 4/27), all represent likely null alleles and three were novel. Compound heterozygosity of a reported nonsense variant and a novel 68-bp deletion was demonstrated for one patient. One patient harbored a reported canonical splice site variant on one allele and a novel frame-shifting duplication on the other allele. Two patients were homozygous for two different nonsense variants, one of them novel.

### RPE65

A total of four novel variants in *RPE65* were identified in two patients (7.4%; 2/27), including one nonsense and three missense. Biallelism could not be formally proven in both cases.

### AIPL1

Of the three variants detected in two affected individuals in *AIPL1* (7.4%; 2/27), there was one novel missense variant found in homozygous state in one patient. Another patient harbored a nonsense variant and a non-canonical splice site change. Whether the variants are in *trans* configuration in this patient could not be established.

### RD3

One patient was found to be homozygous for a known nonsense variant in *RD3* (3.7%; 1/27).

### NMNAT1

*NMNAT1* variants were detected in one patient (3.7%; 1/27) who possessed one reported frame-shifting duplication and the known hypomorphic variant c.769G>A/p.E257K [16]. Biallelism could not be confirmed due to lack of additional family DNA samples.

### CEP290

One patient was found to be homozygous for the common c.2991+1655A>G/p.C998* allele which causes insertion of a cryptic exon and subsequent truncation [5,17].

## Other patients

In addition to the cases described above, we identified eight patients (30%) who harbored pathogenic variants in genes not typically associated with LCA. Clinical re-evaluation of these cases led to a revision of the initial clinical diagnosis in all of them. Within this group, *ABCA4* was the most frequently mutated gene, as biallelic variants were seen in four patients. In these cases, a later onset of disease and a dense pigmentation of the retina were observed (Table 3). After genetic testing and re-evaluation of clinical data, the diagnosis was corrected to cone-rod dystrophy (CRD), demonstrating severe morphological and functional damage in all cases.

In addition, we found a male patient to be hemizygous for a pathogenic variant in *RP2*. He had been initially diagnosed in adult age with severely progressed retinal degeneration. Consequently, his diagnosis was corrected to X-linked retinitis pigmentosa.

Another male patient was shown to be hemizygous for a pathogenic variant in *CACNA1F*. He was suffering from nystagmus, night blindness, photophobia and very poor vision since birth. His fullfield ERGs showed residual photopic and scotopic responses. Morphologically, slight attenuation of the retinal vessels, changes in the macular reflexes and only minimal peripheral pigmentary changes could be observed. In this case, the diagnosis was changed to X-linked congenital stationary night blindness (CSNB).

One patient harbored pathogenic variants in *CDHR1*. The revised clinical diagnosis in this case was CRD, but interestingly, this female patient also suffered from renal insufficiency, secondary hyperparathyroidism and obesity. Whether these symptoms can be considered as a unique disease identity or syndrome remains unexplained. So far, such extra-ocular symptoms have not been described as a feature of *CDHR1*-related disease but would be typical features of a ciliopathy to which *CDHR1*-associated IRD does not belong to.

The last male patient presented in our clinic with a severe retinal degeneration at the age of 50 years and was found to be homozygous for an in-frame insertion/deletion in *PROM1*. On the basis of patient history, clinical findings and genetic results, the clinical diagnosis was changed to autosomal recessive retinitis pigmentosa.

## Discussion

In a cohort of 27 German patients initially diagnosed with LCA, we were able to identify sequence variants likely explaining the disease phenotype in 25 cases (93%) by applying a cost-efficient targeted next-generation sequencing approach designed at the Department of Human Genetics, Radboud University Medical Center, Nijmegen, The Netherlands. The MIP panel targets 108 known IRD genes, including 22 genes that are associated with LCA, that were reported in October 2013.

Undoubtedly, those LCA genes with the highest disease-causing variant load have already been discovered. However, the fact that half of the variants (17/34) we identified are novel suggests that the mutation spectrum of LCA and other IRD genes is far from being saturated and confirms the known genetic heterogeneity of IRD in an outbred European population.

The most frequently mutated LCA genes in our cohort were *CRB1* (6 cases, 22%) and *RPGRIP1* (4 cases, 15%). Among the six patients with *CRB1* mutations, five carried the recurrent p.C948Y variant on one allele, which is known to be a founder mutation [18]. We only identified one patient with a *CEP290* variant in our cohort, despite *CEP290* being one of the most frequently mutated LCA genes in different populations [5, 19], but this is due to the fact that most patients in the present study had already been pre-screened for the recurrent pathogenic intronic variant c.2992+1655A>G.

Several criteria were considered to evaluate the potential pathogenicity of variants: (1) variants have previously been reported to be pathogenic, (2) variants are observed only in few heterozygous cases or are absent among 277,264 general population alleles sourced from gnomAD browser; (3) variants represent likely null alleles (nonsense, canonical splice site and frame-shift variants), and (4) in the case of missense variants they are predicted to be damaging by *in silico* prediction algorithms. In addition, all variants were classified according to their pathogenicity based on the American College of Medical Genetics and Genomics (ACMG) guidelines [8]. With nonsense, canonical splice site and frame-shifting variants having a strong weight in the ACMG scoring system, this class of variants are consequently classified either as likely pathogenic or pathogenic, whereas missense variants that lack segregation data and functional analyses to support a damaging effect are always classified as variants of uncertain significance (VUS). To compensate for this simplistic categorization of the ACMG classification system, we provide *in silico* predictions from four algorithms for all missense variants identified in this study, regardless of having been reported previously or not, along with phyloP scores, Grantham differences and MAFs sourced from the gnomAD browser (Table 2). The extremely low MAF or even the absence in the gnomAD browser, the evolutionary conservation as well as the type of the respective amino acid substitution are strong indicators that all missense variants we identified and reported are indeed pathogenic. One missense variant that is predicted to be benign by the majority of algorithms is the recurrent c.769G>A/p.E257K variant in *NMNAT1*, but it has been shown previously that this is a hypomorphic variant and almost always causes LCA in combination with more severe alleles [16].

Apart from the fact that we lack segregation data for several patients, the only case that is left with some level of uncertainty is patient LCA 108 who carries a nonsense variant and a non-canonical splice site variant in *AIPL1*. The latter is a transition of T to C at position +6 of the splice donor of exon 2. It is absent in the gnomAD browser, but since the +6 position is not invariable, we performed an *in silico* prediction. The bioinformatic tool Human Splicing Finder [20] predicts that the c.277+6T>C variant breaks the natural splice donor site, since the mutant score is reduced by 41% compared to the wildtype score when using maximum entropy as the algorithm type. However, since *AIPL1* is not expressed in accessible tissues like blood or skin fibroblasts, mRNA analyses to confirm the *in silico* prediction are not feasible. Sanger sequencing of the entire coding region of *AIPL1* in this patient revealed no other variants than c.834G>A/p.W278* and c.277+6T>C. While most cases with mutations in *AIPL1* are biallelic, certain mutations may result in dominant cone-rod dystrophy or juvenile retinitis pigmentosa [21], however, this most probably is not the case for loss of function alleles like the c.834G>A/p.W278* variant in our patient. Of course, we cannot rule out that the phenotype of our patient might not be related to *AIPL1* at all.

The different forms of IRD may present with considerable clinical overlap [22]. This often precludes the assessment of a diagnosis on the basis of the disease phenotype alone, no matter how experienced and meticulous the clinician might be. Hence, we were not surprised that eight patients in our cohort (30%) were found to carry pathogenic variants in genes not typically associated with LCA. We reassessed the clinical data of these patients and revisited the initial diagnosis in all of them. A recently published study on Brazilian patients with LCA found the same proportion (i.e. 30%) of patients that were solved by identifying variants in non-LCA genes [23]. This impressively demonstrates how a molecular diagnosis can help to refine a clinical diagnosis.

The underlying variants in two patients remained unresolved (7.4%; 2/27). One of these patients was found to be heterozygous for a known missense variant in *IMPG2*. Biallelic mutations in *IMPG2* are a known cause for RP [24]. All exons and adjacent intronic regions of this gene were sufficiently covered which excludes the existence of a second variant in the coding region. Whether non-coding deep-intronic variants or large deletions in the *IMPG2* gene account for the second pathogenic allele in this patient remains unknown.

Supposing that all patients in whom we could not confirm *trans* configuration of variants are indeed biallelic, our detection rate is 93%. This is in line with recent studies for LCA which achieved 80-90% in panel-based approaches [2-3] and 89% by whole exome/genome sequencing [4]. Analysis of our sequencing data revealed several regions with low or no coverage, as for instance for parts of exon 6 of *CRB1*. We would have missed several patients carrying the recurrent c.2843G>A/p.C948Y variant in this gene, had we not re-sequenced this exon in all patients with conventional Sanger sequencing. It would be interesting to know whether we would have achieved a detection rate of 100% had all genes in our panel been sufficiently covered.

Several studies have shown that whole exome sequencing (WES) and whole genome sequencing (WGS) can outperform targeted sequencing approaches in terms of variant detection [4, 25-27]. In fact, NHS England is already planning to commission WGS into routine clinical care pathways [28]. However, targeted sequencing approaches have several benefits, including a higher coverage rate for targeted regions and higher throughput in terms of patient numbers. What is more important, they are associated with considerable lower costs, which is relevant for those patients who cannot expect reimbursement from their health care provider or have no health insurance at all. The MIP technology we used can be as low as € 80 per sample per gene panel, which is 10 to 20 times lower than the price tag for other NGS-based sequencing procedures. Reaching a detection rate of 93%, we could demonstrate that MIPs are a cost-efficient and sensitive tool for targeted next-generation sequencing in IRD.

## Acknowledgements

This work was supported by grants from the Pro Retina Germany Foundation (Pro-Re/Project/Zobor.1-2016) ot NW, BF, SK and DZ and by the Curing Retinal Blindness Foundation and the Candle in the Dark-Childvision Research Fund, managed by the King Baudouin Foundation to FPMC and MIK. We further acknowledge support by the Deutsche Forschungsgemeinschaft and the Open Access Publishing Fund of the University of Tuebingen.

## References

1. Chung DC, Traboulsi EI. Leber congenital amaurosis: clinical correlations with genotypes, gene therapy trials update, and future directions. J AAPOS. 2009;13: 587–92.

2. Bernardis I, Chiesi L, Tenedini E, Artuso L, Percesepe A, Artusi V et al. Unravelling the Complexity of Inherited Retinal Dystrophies Molecular Testing: Added Value of Targeted Next-Generation Sequencing. Biomed Res Int. 2016;2016: 6341870.

3. Thompson JA, De Roach JN, McLaren TL, Montgomery HE, Hoffmann LH, Campbell IR et al. The genetic profile of Leber congenital amaurosis in an Australian cohort. Mol Genet Genomic Med. 2017;5: 652–67.

4. Carss KJ, Arno G, Erwood M, Stephens J, Sanchis-Juan A, Hull S et al. Comprehensive Rare Variant Analysis via Whole-Genome Sequencing to Determine the Molecular Pathology of Inherited Retinal Disease. Am J Hum Genet. 2017;100: 75–90.

5. den Hollander AI, Koenekoop RK, Yzer S, Lopez I, Arends ML, Voesenek KE et al. Mutations in the CEP290 (NPHP6) gene are a frequent cause of Leber congenital amaurosis. Am J Hum Genet. 2006;79: 556–61.

6. Li H, Durbin R. Fast and accurate short read alignment with Burrows-Wheeler transform. Bioinformatics. 2009;25: 1754–60.

7. Lek M, Karczewski KJ, Minikel EV, Samocha KE, Banks E, Fennell T et al. Exome Aggregation Consortium. Analysis of protein-coding genetic variation in 60,706 humans. Nature. 2016;536: 285–91.

8. Richards S, Aziz N, Bale S, Bick D, Das S, Gastier-Foster J et al. ACMG Laboratory Quality Assurance Committee. Standards and guidelines for the interpretation of sequence variants: a joint consensus recommendation of the American College of Medical Genetics and Genomics and the Association for Molecular Pathology. Genet Med 2015;17: 405–24.

9. Kumar P, Henikoff S, Ng PC. Predicting the effects of coding non-synonymous variants on protein function using the SIFT algorithm. Nat Protoc 2009; 4: 1073−81.

10. Adzhubei IA, Schmidt S, Peshkin L, Ramensky VE, Gerasimova A, Bork P et al. A method and server for predicting damaging missense mutations. Nat Methods 2010;7: 248−249.

11. Schwarz JM, Cooper DN, Schuelke M, Seelow D. MutationTaster2: mutation prediction for the deep-sequencing age. Nat Methods 2014;11: 361−362.

12. Choi Y, Chan AP. PROVEAN web server: a tool to predict the functional effect of amino acid substitutions and indels. Bioinformatics. 2015;31:2745–7.

13. Landrum MJ, Lee JM, Riley GR, Jang W, Rubinstein WS, Church DM et al. ClinVar: Public archive of relationships among sequence variation and human phenotype. Nucleic Acids Res. 2014;42(Database issue): D980–5.

14. Hanein S, Perrault I, Gerber S, Tanguy G, Barbet F, Ducroq D et al. Leber congenital amaurosis: comprehensive survey of the genetic heterogeneity, refinement of the clinical definition, and genotype-phenotype correlations as a strategy for molecular diagnosis. Hum Mutat. 2004;23: 306–17.

15. Corton M, Tatu SD, Avila-Fernandez A, Vallespín E, Tapias I, Cantalapiedra D et al. High frequency of CRB1 mutations as cause of Early-Onset Retinal Dystrophies in the Spanish population. Orphanet J Rare Dis. 2013;8: 20.

16. Siemiatkowska AM, van den Born LI, van Genderen MM, Bertelsen M, Zobor D, Rohrschneider K et al. Novel compound heterozygous NMNAT1 variants associated with Leber congenital amaurosis. Mol Vis. 2014;20: 753–9.

17. Parfitt DA, Lane A, Ramsden CM, Carr AJ, Munro PM, Jovanovic K et al. Identification and Correction of Mechanisms Underlying Inherited Blindness in Human iPSC-Derived Optic Cups. Cell Stem Cell. 2016;18: 769–81

18. den Hollander AI, Roepman R, Koenekoop RK, Cremers FP. Leber congenital amaurosis: genes, proteins and disease mechanisms. Prog Retin Eye Res. 2008;27: 391–419.

19. Coppieters F, Casteels I, Meire F, De Jaegere S, Hooghe S, van Regemorter N et al. Genetic screening of LCA in Belgium: predominance of CEP290 and identification of potential modifier alleles in AHI1 of CEP290-related phenotypes. Hum Mutat. 2010;31: E1709–66.

20. Desmet FO, Hamroun D, Lalande M, Collod-Béroud G, Claustres M, Béroud C. Human Splicing Finder: an online bioinformatics tool to predict splicing signals. Nucleic Acids Res. 2009;37: e67.

21. Sohocki MM, Perrault I, Leroy BP, Payne AM, Dharmaraj S, Bhattacharya SS et al. Prevalence of AIPL1 mutations in inherited retinal degenerative disease. Mol Genet Metab. 2000;70: 142–50.

22. Sahel JA, Marazova K, Audo I. Clinical characteristics and current therapies for inherited retinal degenerations. Cold Spring Harb Perspect Med 2014;5: a017111.

23. Porto FBO, Jones EM, Branch J, Soens ZT, Maia IM, Sena IFG et al. Molecular Screening of 43 Brazilian Families Diagnosed with Leber Congenital Amaurosis or Early-Onset Severe Retinal Dystrophy. Genes (Basel). 2017;8(12). pii: E355.

24. Bandah-Rozenfeld D, Collin RW, Banin E, van den Born LI, Coene KL, Siemiatkowska AM et al. Mutations in IMPG2, encoding interphotoreceptor matrix proteoglycan 2, cause autosomal-recessive retinitis pigmentosa. Am J Hum Genet. 2010;87: 199–208.

25. Lelieveld SH, Spielmann M, Mundlos S, Veltman JA, Gilissen C. Comparison of Exome and Genome Sequencing Technologies for the Complete Capture of Protein-Coding Regions. Hum Mutat 2015;36: 815–22.

26. Meienberg J, Bruggmann R, Oexle K, Matyas G. Clinical sequencing: is WGS the better WES? Hum Genet 2016;135: 359–62.

27. Ellingford JM, Barton S, Bhaskar S, Williams SG, Sergouniotis PI, O’Sullivan J et al. Whole Genome Sequencing Increases Molecular Diagnostic Yield Compared with Current Diagnostic Testing for Inherited Retinal Disease. Ophthalmology 2016;123: 1143–50.

28. Turnbull C, Scott RH, Thomas E, Jones L, Murugaesu N, Pretty FB et al. The 100 000 Genomes Project: bringing whole genome sequencing to the NHS. BMJ. 2018;361: k1687.

